# Echo state networks with multiple readout modules

**DOI:** 10.1101/017558

**Authors:** Andres Laan, Raul Vicente

**Author notes:** Also at Champalimaud Neuroscience Programme, Champalimaud Foundation, Avenida Brasilia, 1400-038 Lisbon, Portugal (Andres Laan).

## Abstract

We propose a new readout architecture for echo state networks where multiple linear readout modules are activated at distinct time points to varying degrees by a separate controller module. The controller module, like the reservoir of the echo state network, can be initialized randomly. All linear readout modules are trained through simple linear regression, which is the only adaptive step in the modified algorithm. The resulting architecture provides modest improvements on a variety of time series processing tasks (between 5 to 50% in performance metric depending on the task studied). The novel architecture is guaranteed to perform at least as accurately as a conventional linear readout. It can be utilized as a general purpose readout method when augmentations to performance relative to the standard method is needed.

**PACS numbers:** 05.45.Tp, 07.05.Mh

## I. INTRODUCTION

Function approximation methods seek to learn mappings from the input feature space *x* to the output feature space *y*. In parametric methods, a general mapping function is used which can approximate a wide variety of different functions depending on the precise value of its parameters. The values of the parameters are learned from example data. Perhaps the simplest parametric mapping function is linear regression, where the output is expressed as a linear combination of input signals. When linear regression fails to provide adequate precision, a common remedy involves calculating a nonlinear expansion *f* (*x*) of the input feature space and performing linear regression between *f* (*x*) and *y* [1]. For time series analysis, echo state networks (ESNs) provide one such expansion function [2]. ESNs have been successfully applied to a wide variety of time series processing problems such as chaotic time series prediction, nonlinear system identification and classification.

In general, the precise conditions on the true mapping function from *x* to *y* for which a linear readout of the nonlinear expansion is sufficient are not known for many expansion methods (but see [3]). We first present a heuristic argument that a single linear readout is expected to be insufficient for some classes of data streams. Then, we propose an elaboration of the basic ESN design that provides greater expressive power than a single linear readout while retaining the property that the only adaptive step in the training process involves simple linear regression.

### A. Motivation

For concreteness, we consider the task of predicting the future behavior of an animal. For many simple animals such as worms, flies and fish, a large fraction of their time is spent performing stereotypical behavioral programs such as crawling, grooming, walking, eating, just to mention a few examples [4]. Every class of stereotyped behavior is conceptualized as motion along a lowdimensional manifold (typically 1D in the intrinsic coordinate system). Switching between programs corresponds to a switch from one manifold to another. The behavior of the animal can be described in terms of the magnitudes of its intrinsic coordinates (muscle activities) or some more easily observed proxies such as joint angles or body postures. For motion along some one-dimensional manifold, any intrinsic coordinate x evolves according to *x* = *X* (*t*), where *t* represents both time and the natural manifold coordinate. Predicting future behavior *dt* time steps ahead requires finding a mapping from *X*(*t*) to *X*(*t* + *dt*). By the inverse function theorem *t* is also a function of *x* (*t* = *X*^−1^(*x*)) in the neighborhood of any point of its domain and we can express *t* as a Taylor series in *x*:

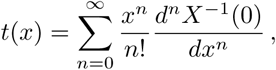

where *X*^−1^ is the inverse function from *x* to *t*. As *x* is also a Taylor series of *t*:

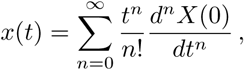

we can write *x*(*t* + *dt*) as a Taylor series of *x*:

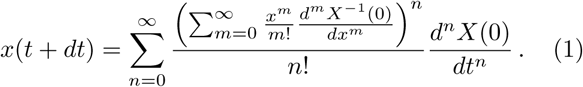

Such Taylor corresponds to a linear readout of a polynomial expansion of *x*.

When motion of the animal switches from one manifold to another, the predictive linear readout of the expansion generally changes, because the functional form of the derivatives (which are directly related to the values of the linear readout coefficients) also change. Thus, for perfect prediction of the trajectories of an object whose motion switches between manifolds, different linear readouts at different times are required.

## II. THE ARCHITECTURE

Extending the polynomial expansion analogy to echo state networks, we might expect mappings of complicated dynamical systems to be better approximated by a combination of linear readouts used to different extents at different times. If the reservoir state at time *t* is row vector *R*(*t*), we can write the predicted output *y_p_*(*t*) as a linear sum of the predictions of individual readout modules *y_p_*(*t*) = ∑*_i_P_i_*(*t*)*y_pi_*(*t*) where *p_i_*(*t*) is the time dependent module weight which lies between 0 and 1, and *y_pi_*(*t*) represents the prediction made by module *i* at time *t*. We restrict ourselves to a scalar output *y_p_*(*t*) for notational clarity, the generalization to a vector output is straightforward.

For each module prediction *y_pi_*(*t*) = *R*(*t*)*W_i_*, where *W_i_* is the column vector of linear readout weights for module *i*. The weights *p_i_*(*t*) are calculated by a separate softmax control module: *p_i_*(*t*) = *exp*(*R*(*t*) ∗ *w_i_*)/∑ *exp*(*R*(*t*) ∗ *w_j_*). The lowercase *w_i_* denote the column vector of weights for the softmax of each module. For a suichoice of *w_i_*, each module *i* will dominate the predicted output *y_p_*(*t*) for those time periods where *p_i_*(*t*) has a high value. This allows the individual readout modules *W_i_* to de facto specialize on predicting the outputs at certain epochs, while their possibly erroneous contributions will be ignored for other epochs where their *p_i_* (*t*) acquires low values. See Fig. 1 for a schematic of the architecture.

**FIG. 1.**
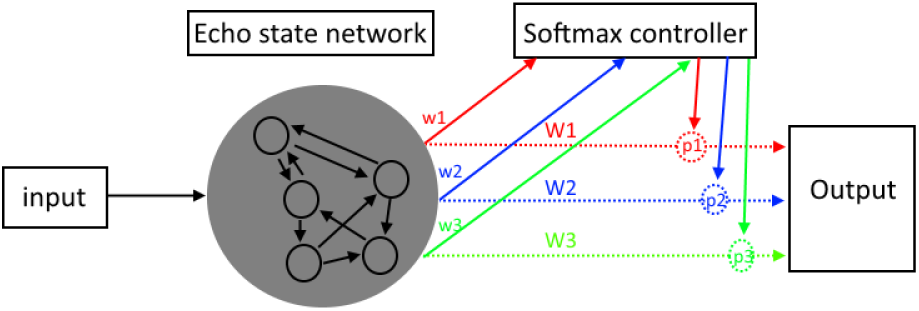
Schematic of the proposed architecture. Multiple linear readouts of a single ESN are adaptively activated to varying degrees by a separate controller module.

The overall aim of the architecture is to reduce the squared error between the desired *y* and the predicted output 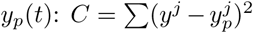, where the sum runs over all the training data examples. The weights *w_i_* and *W_i_* can in principle be trained by joint gradient descent to reduce the cost function. However, a simpler approach works just as well in practice where in *w_i_* are initialized randomly and all the *W_i_* are trained jointly as a least squares problem.

If we fix the *w_i_*, minimizing the cost function can be converted to an ordinary least square regression problem. The column vector *y_p_*(*t*) can be expressed as the matrix product *R_p_W*, where *W* is the columnwise concatenation of the individual readout modules 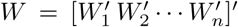. Row *t* of the matrix *R_p_* corresponds to a row-wise concatenation of vectors *p_i_*(*t*)*R*(*t*): *R_p_*(*t*) = [*p*_1_(*t*)*R*(*t*)*p*_2_(*t*)*R*(*t*) ⋯ *p_n_*(*t*)*R*(*t*)]. The *p_i_*(*t*) needed to form the matrix *R_p_*(*t*) are calculated based on the fixed *w_i_* and reservoir neuron activities. The soft-max weights *w_i_* are initialized randomly from a uniform distribution between −1 and 1 and then multiplied by a constant. A good choice of the constant is critical for effective results. For low values of the constant, all weights *p_i_*(*t*) remain near the value 1/*n* during all time points, thus preventing the readout modules from specializing. For very high values of the constant, the weights *p_i_*(*t*) jump rapidly between 0 and 1, creating a lack of smoothness. For a suiinitialization, where weights *p_i_*(*t*) smoothly vary over time between values 0.1 and 0.9, the network performance is very close to the best results achieved by gradient descent, while the time required to estimate the final *W_i_* is dramatically shortened. Finally, we note that the network is guaranteed to work as well as a single module readout. If all *W_i_* are set to equal the single module readout *W_s_*, then the prediction produced by the new network is equivalent to the prediction produced by *W_s_*, because weights *p_i_*(*t*) sum to unity for each time step.

## III. RESULTS

We first tested the performance of the new architecture on a behavior prediction task. We created a simulated lamprey, which switches between different behavior modes of swimming, digging and struggling. The lamprey was chosen because its multisegment body shows remarkably reliable behavior well-described by simple mathematical equations [5–7]. The body of the model lamprey consisted of 20 segments. The equation describing swimming was written as *a_i_* = *sin*(2*πi*/*l* − 2*πt*/*T*), where *i* is the segment number, *a_i_* is the activity of segment *i* and *l* and 1/*T* are the wavelength and frequency of the wave respectively. Like in the real lamprey, the wavelength was kept fixed while the period *T* varied between 20 and 200. The equation for digging was *a_i_* = *sin*(2*πi*/40)*sin*(2*πft*), where *f* varied from 0.05 and 0.0125 cycles/s. In the real lamprey, struggling consists of a periodic contortion of the body into a convoluted shape, but is less well understood mathematically. We choose to model struggling using the equation *a_i_* = *sin*(2*πi* − 10)/(20 + 10*sin*(2*πft*)); see Figure 2 for a graphical summary of all modes. A small amount of uniform random noise with amplitude 0.1 was added to each *a_i_* at all time steps.

**FIG. 2.**
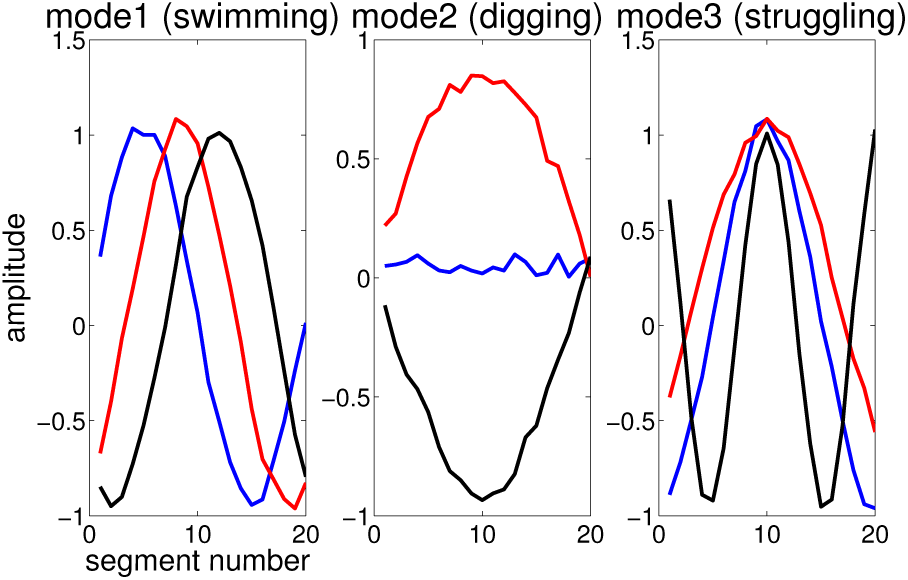
Illustration of the three modes of worm behavior-swimming (a translated sinusoid), digging (a vibrating string) and struggling (a contracting deformation of the body). Different lines in a given mode correspond to different snapshots of the amplitude of the different body segments. The delay between the blue and red and the red and black traces is seven time steps. In the prediction task, the prediction horizon was 10 time steps.

Each behavioral mode had a duration of 300 time steps. After the completion of 300 time steps, the lamprey switched between the modes randomly. For each 300 step epoch, the equation control parameters 1/T or f remained fixed, but between epochs the parameters switched randomly. The task of the echo state network was to predict the body posture of the model lamprey 10 time steps ahead of the present posture of the lamprey. For this task the network was trained with a randomly initialized time series of 10000 time steps and test error was measured on a differently initialized time series of equal length, and a linear model was fitted to relate *R*(*t*) (calculated at each time step using *R*(*t* − 1) and all the *a_i_*(*t*)) to each *a_i_*(*t* + 10). For a reservoir with 100 neurons (connection probability 0.05, largest connectivity matrix eigenvalue 0.95), which was augmented with a further set of 100 features found by calculating the square of each reservoir neuron activation at every time step [8], the echo state network achieved a normalized mean square prediction error 0.360+/−0.014 (for worm motion prediction all nmse reported as the averages of 50 random reservoir initializations +/−s.e.m), while a 2-module readout give an improvement of 30% (nmse 0.25+/−0.0065) and a 3-module readout give an improvement of 35% (0.234+/−0.007) compared to the single module prediction.

A 3-module readout has 600*20 adjustable free parameters compared to the 200*20 free parameters of a single module architecture. Another possible comparison can thus be made to a 300 neuron reservoir with quadratic feature augmentation, a model which also has 600*20 adjustable parameters. The mean test set nmse on 50 trials (both input and reservoirs randomly reinitialized at each trial) gave a prediction improvement of 35% (0.233+/−0.009) relative to the 100 neuron single-readout reservoir-a result statistically indistinguishable from the improvement given by the 3-module readout with a 100 neuron reservoir. However, the 100 neuron reservoir with 3 readout modules has a considerably smaller run-time complexity. To get the next reservoir state, 300*300 multiplications must be carried out at run time for 300 neuron reservoir compared to the 100*100 multiplications that must be carried out for use of the 100 neuron 3-module method.

Three methods (feature augmentation with polynomials of reservoir activities, increasing the reservoir size or introducing multiple readout modules) give complementary improvements to each other. For equivalent number of introduced adjustable parameters, they provide comparable improvements to performance. For another example, a 100 neuron reservoir without feature augmentations gives a test-set nmse of 0.44+/−0.0095, which is improved by quadratic feature augmentation to 0.36, while a 2-module readout (without quadratic augmentation) gives an nmse of 0.38+/−0.015 and a 200-neuron reservoir with a single readout module and no quadratic augmentation gives an nmse of 0.35+/−0.012. Conceptually, the three methods of improving ESN performance provide different benefits. While they all increase the number of adjustable parameters, larger reservoirs provide a larger working memory [9], while multi-module readouts allow prediction modules to fine tune their predictions for distinct phases of behavior. Both quadratic augmentation and multiple readout modules provide improved performance with only marginal increases in model runtime complexity, especially when compared to increases in reservoir size.

Next, we tested the new architecture on the standard Mackey-Glass (MG) chaotic time-series prediction task [10]. For MG(17) with reservoir size 1000, no significant improvements were found when comparing a 2-module method with the standard single module method. When the reservoir size was reduced to 500, the 2-module method gave significant improvements. We measure the performance of the method by calculating the time when the difference between the predicted and actual continuation became larger than 0.1, which we call the divergence time. The divergence time rose from 990 to 1500 when comparing the 1-module case with the 2-module system. Also, the standard deviation of the divergence time decreased significantly from 500 to 350 for the 2-module readout, this despite an increase of the divergence time itself for the 2-module case. A re-implementation of Jaeger 2004 (where ESN prediction of the MG system was first reported) with 1000 reservoir neurons found divergence time of around 1640 with a standard deviation of 400. Thus, a 2-module readout method is able to achieve performance that is only 10% inferior to a 1000 neuron reservoir but has a 4 times lower run-time complexity.

**FIG. 3.**
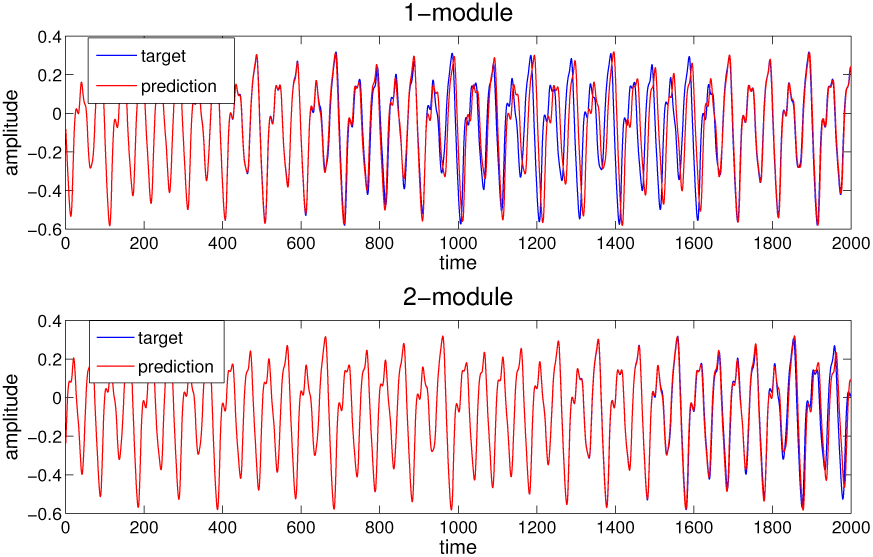
Example of performance on the Mackey-Glass time series prediction (blue traces) for a 500 neuron reservoir with 1 and 2 readout modules, respectively. The 2-module readout system clearly tracks the correct continuation (red traces) for a longer duration. The zero time point marks the end of training and the beginning of the prediction phase.

**FIG. 4.**
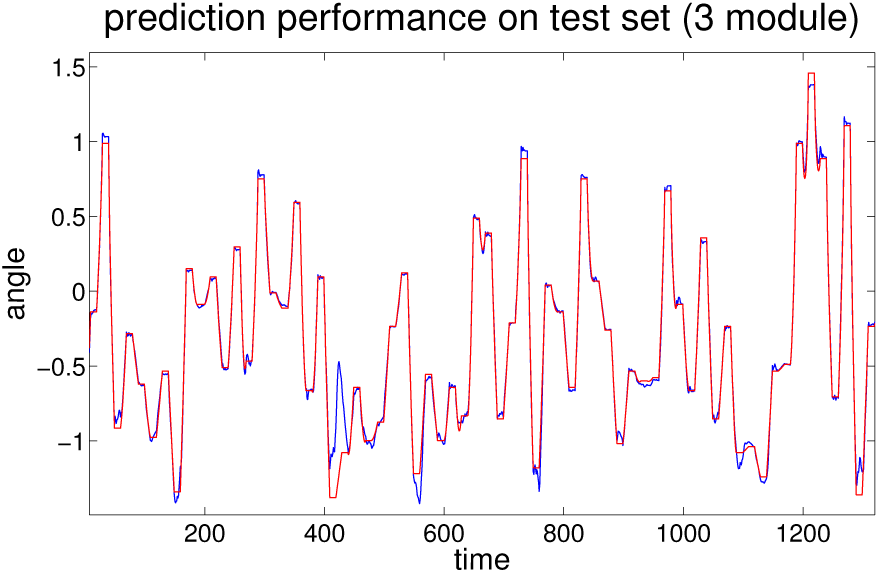
Prediction (blue) versus actual signal shown for test data for a 3-module readout system in the coordinate trans-formation task.

Another time series prediction problem involves forecasting electricity loads. We obtained freely available data from the 2012 Global Energy Forecasting Competition on the data science competitions website *kaggle.com.* The dataset contained 20 timeseries of hourly loads for 20 different utility zones spanning a period of four years. Hourly temperature data was available for 11 stations as well. We used ESNs to predict the summed load of the 20 utility stations using as inputs the time of day, the month, and temperature signals. Table 1 shows the test set nmse values (average of 50 reservoir initializations) for a reservoirs with 100, 200 and 400 neurons and compares these to a 2-module readout and quadratic augmentation for N=100 and N=200 neuron reservoirs. For this dataset-comparing methods with equal numbers of free parameters-quadratic augmentation gives the largest improvements, followed by the 2-module architecture. Increasing the reservoir size performs worst in this dataset.

**TABLE I.** Results for different methods.

| Method | Reservoir size | Number params. | NMSE | S.D. |
| --- | --- | --- | --- | --- |
| Standard | 100 | 100 | 0.168 | 0.0043 |
| Standard | 200 | 200 | 0.1526 | 0.0065 |
| Standard | 400 | 400 | 0.1373 | 0.0067 |
| Quad. augment. | 100 | 200 | 0.111 | 0.0041 |
| Quad. augment. | 200 | 400 | 0.107 | 0.003 |
| 2-module | 100 | 200 | 0.1365 | 0.01 |
| 2-module | 200 | 400 | 0.125 | 0.009 |

On the Santa Fe time series prediction task D, the 2-module readout gave a modest 5% improvement over the 1-module method for prediction delays between 2 to 5 time steps.

Finally, we tested the new architecture on a motor coordinate transformation task. Here, a model human with two joints had to move his arm between randomly chosen points on the (x,y) plane. A sequence of 500 points were chosen randomly within a circular quadrant that was within reach of the model arm. The task was to move the arm with uniform speed from one point to the next in 10 time steps, then hold the hand steady for 10 steps until moving forward to the next point. The input positions were specified in normalized x,y coordinates normalized to the 0 to 1 range, whereas the output was specified in radian joint angle coordinates from which 1.4 was subtracted to remove most of the mean. Performance of a single module ESN with 500 neurons gave a small NMSE of 0.017. A 3-module ESN improved performance by 50%. Note that in this task the inertia of the model arm was not modeled, so the task could in principle have been solved by a feed forward, not a recurrent neural net. It was studied primarily because its sequences of dynamics and stability might pose problems for an echo state network whose neurons can show ongoing dynamics also during stable phases of the task.

## IV. DISCUSSION

Many simple tricks have been proposed to augment the performance of ESNs. These include increasing nonlinearity by augmenting the non-linear expansion with polynomial functions of reservoir activities [8], increasing the reservoir size [10], averaging predictions from many reservoirs [10], introducing delay lines into the read out system [11], providing neurons with a diversity of time constants and having the reservoir adapt to input statistics via intrinsic plasticity [12]. The new multi-module readout architecture proposed is in its simplest form equivalent to introducing an additional layer of fixed non-linearity into the readout layer for improved performance. However, in principle, the new layer of nonlinearity is trainable and it might still be the case that for certain tasks, gradient descent of the softmax weights *w_i_* produces far superior results to the random initialization. In its randomly initialized form, it is most likely useful as an out-of-the-box non-linearity which can be tried when the traditional tricks run short of providing the required performance. It is guaranteed to work as well as a single readout module or better with very little additional training cost. We have demonstrated its modest usefulness on a variety of data series processing tasks. We also note that the idea of training a controller for multiple ESN readouts can be understood as an adaptation of ensemble methods to reservoir computing. Future work will focus on robust architectures for learning while coping with slowly-varying non-stationary signals.

## ACKNOWLEDGEMENTS

We thank the financial support of the Estonian Research Council via the grant PUT438.

